# Solid immersion microscopy readily and inexpensively enables 12 nm resolution on plunge-frozen cells

**DOI:** 10.1101/373647

**Authors:** Lin Wang, Benji Bateman, Laura C. Zanetti-Domingues, Amy N. Moores, Sam Astbury, Christopher Spindloe, Michele C. Darrow, Maria Romano, Sarah R. Needham, Konstantinos Beis, Daniel J. Rolfe, David T. Clarke, Marisa L. Martin-Fernandez

**Affiliations:** Science and Technology Facilities Council, Central Laser Facility, Rutherford Appleton Laboratory, Didcot OX11 0QX, United Kingdom; Diamond Light Source, Harwell Science and Innovation Campus, Didcot OX11 0DE, United Kingdom; Department of Life Sciences, Imperial College London, London SW7 2AZ, United Kingdom; Research Complex at Harwell, Rutherford Appleton Laboratory, Didcot OX11 0FA, United Kingdom

**Keywords:** Cryo-fluorescence microscopy, solid immersion lens, STORM, super-resolution, cell imaging

## Abstract

Super-resolution fluorescence microscopy achieves 20-30 nm resolution by using liquid-immersion objectives to optimize light collection and chemical sample fixation to minimize image blurring. It is known that fluorophore brightness increases substantially under cryogenic conditions and that cryo-fixation is far superior in preserving ultrastructure. However, cryogenic conditions have not been exploited to improve resolution or sample quality because liquid immersion media freezes at the objective, losing its optical properties. Here, simply by replacing the immersion fluid with a low-cost super-hemispherical solid immersion lens (*super*SIL), we effortlessly achieve <8 nm localisation precision and 12 nm resolution under cryogenic conditions in a low-cost, low-tech system. This is to our knowledge the best resolution yet attained in biological samples. Furthermore, we demonstrate multicolour imaging and show that the inexpensive setup outperforms 10-fold more costly super-resolution microscopes. By also removing the barrier to total internal reflection fluorescence imaging of mammalian cells under cryogenic conditions, s*uper*SIL microscopy delivers a straightforward route to achieve unmatched nanoscale resolution on both bacterial and mammalian cell samples, which any laboratory can effortlessly and inexpensively implement.

## Introduction

Super-resolution fluorescence microscopy (FM) techniques, such as structured illumination microscopy (SIM) [1], stimulated emission depletion (STED) microscopy [2], and single molecule localization microscopy (SMLM) (including stochastic optical reconstruction microscopy (STORM) [3], and photoactivated localization microscopy (PALM) [4, 5]), have provided nanoscale insights into the functioning of cellular components for over a decade. However, technical complexity and high costs have so far prevented their general application to the biomedical field outside specialist laboratories. Furthermore, nanoscale resolution typically requires chemical fixation to prevent blurring. In electron microscopy (EM), vitrification through rapid freezing has proven more effective at preserving ultrastructure [6], yet there are few examples in the literature of its use in super-resolution. This is despite the resolution improvement that can be expected from the increased photon budget available under cryogenic conditions. To achieve super-resolution in cryogenic conditions is a timely goal to realize the promise of correlative light and electron microscopy (CLEM) in biology [7], so far limited by the resolution mismatch between cryo-EM and cryo-FM, largely restricting the optical element to identification of regions of interest.

The resolution of SMLM techniques, like STORM, depends on the precision by which individual molecules can be localized in the sample. This in turn depends on the number of photons emitted, which increases under cryogenic conditions, and the number collected by the objective lens, which is determined by its numerical aperture (NA) [8, 9]. NAs > 1 and preferably > 1.4 are required to achieve high resolution, but these need immersion fluids to couple the sample to the objective. Because liquid media freeze, dry objectives (NA ≤ 0.9) are mostly used under cryogenic conditions, with the disadvantages of lower resolution and limited photon collecting ability. The larger photon yield of common fluorophores under cryogenic conditions nevertheless allowed Kaufman *et al* [10] to achieve ~125 nm resolution on green fluorescence protein (GFP)-stained samples. Liu *et al* [11] attained ~74 nm resolution using DRONPA, a protein 2.5 times brighter than GFP. Recent work [12, 13] reported highly specialized cryo-STORM systems functioning at liquid Helium temperature, which yielded an astonishing ~Angstrom localization precision on isolated molecules. Other set-ups relied on custom-built stages to incorporate cryofluids; Nahmani *et al* [14] achieved ~35 nm resolution using a water-immersion objective (NA = ~1.2). A similar approach was taken by Faoro *et al* [15], but this set-up was not applied to super-resolution microscopy. However, the complex machinery required has rendered cryo-STORM so far largely inapplicable to general cell biology. Furthermore, lenses with NAs <1.4, which cannot create evanescent fields, are incompatible with the TIRF illumination required for super-resolution imaging of mammalian cells [16].

Super-hemispherical solid immersion lenses (*super*SILs) are truncated balls made of solid materials of high refractive index and fill the gap between the objective and the sample, eliminating the requirement for coupling fluids [17-19]. By using a *super*SIL to couple the sample to a dry objective, the effective NA of the latter is enhanced up to the value of the refractive index of the *super*SIL. We demonstrated the high-resolution of *super*SIL-based FM under low signal-to-noise-ratio conditions at room temperature [20, 21]. Proof-of-concept experiments were also carried out in STED [22, 23] and SIM [24]. However, *super*SILs are particularly suited to STORM techniques because of the critical dependence on high photon collection efficiency. Indeed, it was previously shown that *super*SILs increased photon collection efficiency by ~6-fold [25].

Here, we describe a low-cost, low-complexity STORM setup that exploits solid immersion technology to effortlessly achieve an unprecedented 12 nm resolution. We solved the problem of how to exploit the increased fluorophore brightness under cryogenic conditions and removed the barrier to TIRF imaging of cryo-vitrified samples and outperformed a much more expensive state-of-the art system at room temperature too. We expect these properties to make *super*SIL microscopy the method of choice for the general implementation of nanoscale imaging in non-specialist laboratories.

## Results and Discussion

Our aim was to find a low-tech solution that could be effortlessly and inexpensively implemented in conventional low-cost optical microscopes and had the flexibility to exploit cryogenic conditions when improving resolution and/or avoiding artefacts from chemical fixation were necessary (**Fig. 1**). The optics to deliver and collect light consist of a 0.55 NA 100× dry objective (Mitutoyo 100× Plan Apo SL Infinity Corrected) and a 1 mm diameter *super*SIL made of cubic zirconia (effective NA = 2.17). An achromatic doublet lens with a focal length of 200 mm was used as tube lens before the EMCCD detector camera (Andor, iXon+ DU-897). The *super*SIL was mounted into the central hole of a platinum disk using a thermally conductive cryo-adhesive (Loctite Stycast 2850 FT epoxy encapsulant) (**Fig. 1A**). *Super*SIL assemblies are robust and inexpensive (~$20), and can be reused by cleaning the surfaces with trypsin, common solvents, and/or piranha solution.

**Fig. 1.**
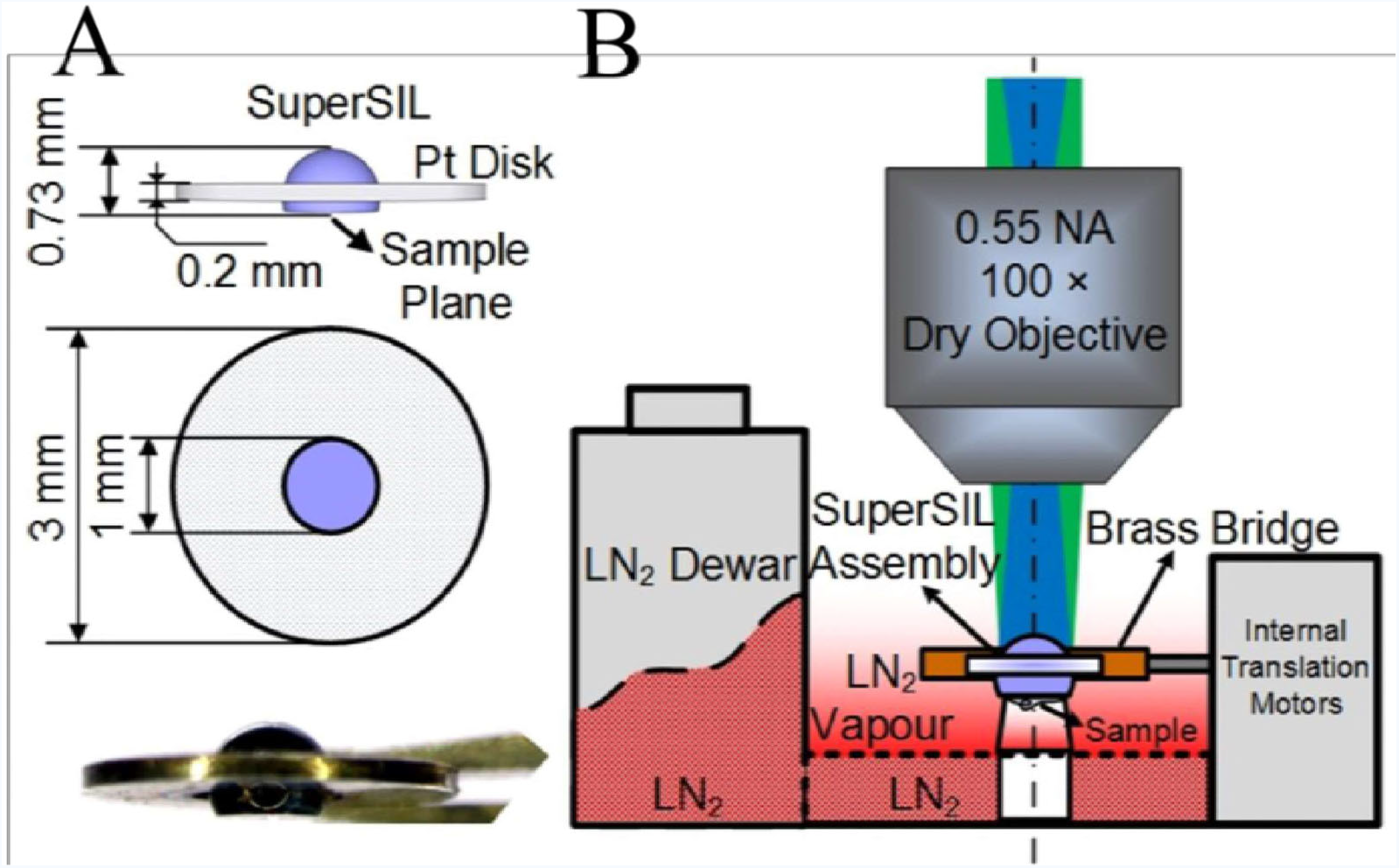
Schematic of *super*SIL microscope. **(A)** Side and top view of *super*SIL assembly. The side view shows the location of the sample plane (aplanatic surface). The photo of an assembly is at the bottom of the panel. **(B)** Schematic of key components in the microscope. An upright microscope configuration was employed. The blue shading illustrates the Köhler illumination laser beam, the green shading indicates the fluorescence emission. (For more info, see **Supporting Information, Fig. S1-S3**).

The *super*SIL assembly was designed to be compatible with the standard EM grid-holder of an off-the-shelf liquid nitrogen (LN_2_)-cooled cryo-stage (Linkam, CMS-196). The samples were adhered to the flat surface of the *super*SILs and plunge-frozen together in liquid ethane. The convex surfaces of the *super*SILs faced upwards to the objective, and the flat surfaces, onto which the samples are adhered, faced downwards (**Fig. 1B**). *Super*SIL assemblies thus play at any temperature, including cryogenic conditions, the role played by liquid immersion media at room temperature. The *super*SIL assemblies were mounted on top of the brass bridge in the cryo-stage and translated in XY directions for fine positioning. The cryo-stage is self-contained with a built-in LN_2_ reservoir, and can be easily integrated into any conventional upright fluorescence microscope. Frozen samples were kept at - 196 °C by the LN_2_ vapour surrounding the brass bridge of the cryo-stage.

The first step to characterize the performance of the *super*SIL microscope was to measure the size of the point spread function (PSF) of the microscope, i.e. the image of a point object, at room temperature, for which we did not cool the stage. We acquired 500 photoluminescence images of the sparse cubic zirconia defect spots present on the flat surface of the *super*SILs (**Fig. S4**), and measured their full width at half maximum (FWHM). These intrinsic defects are single point emitters [26], and are located at the aplanatic plane of the *super*SIL, which is the position with the fewest optical aberrations [20]. Thus, their images give an estimation of the best lateral resolution [27]. Representative images and the statistical results show a resolution of 153 ± 15 nm (**Fig. 2A)**, in agreement with that previously reported by a similar system [21]. The ~25% discrepancy between experimental and theoretical values (118 nm predicted using the Houston criterion [28]) arises from the failure to collect rays propagating at angles approaching 90° [20].

**Fig. 2.**
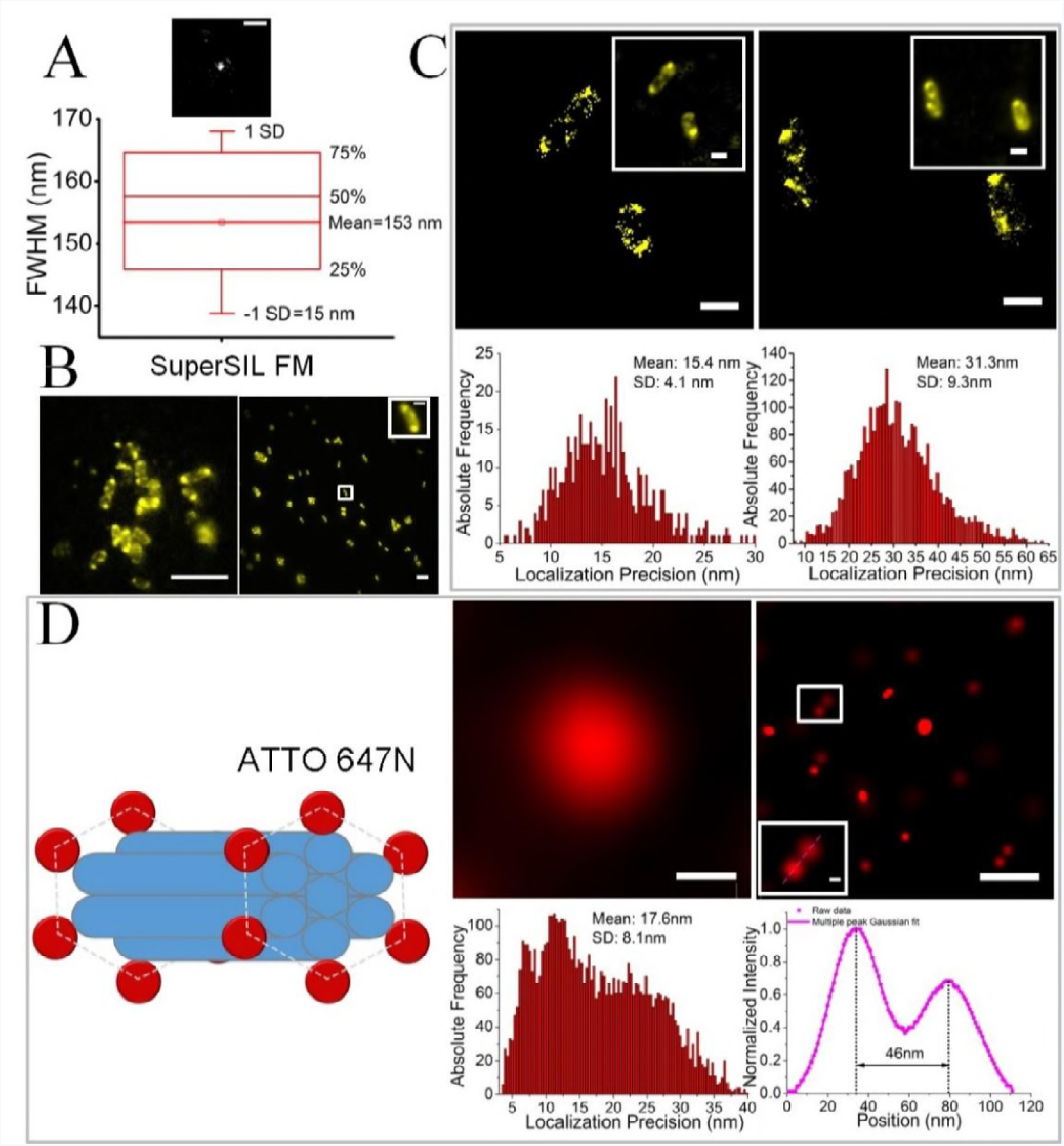
S*uper*SIL resolution performance at room temperature. **(A)** Image of a point object (*top*). Bar: 0.5 μm. The box chart (*bottom*) of the full width half maximum (FWHM) measurements of images from point emitters. The boxes show the 25th and 75th percentiles of the data points; the whiskers the standard deviation. **(B)** Comparison of wide-field images of live McjD-EGFP in *E.coli* cells. The *super*SIL image (*left*) has 471× magnification, while the specialist microscope (*right*) has 100× magnification. For comparison the inset contains a 4.71× scaled-up image of a cell. Bar: 5 μm, and 1 μm in the inset. **(C)** STORM images (*top*) and precision histograms (*bottom*) of live McjD-EGFP *E.coli* cells in *super*SIL STORM (*left*) and specialist STORM (*right*). The insets show wide-field fluorescence images of the cells. Bar: 1 μm. **(D)** Resolution evaluation of STORM imaging in the specialist microscope. *Left*: Schematic of DNA origami nanorulers labelled with ATTO 647N dye molecules. *Right*: image of a field of nanorulers obtained from wide-field (*top left*) and STORM (*top right*). Bar: 200 nm. The STORM image of a nanoruler is shown in the inset, indicated by the white-border box. Bar: 20 nm. *Bottom left*: Localization precision histogram from the STORM image of the nanorulers. *Bottom right*: Line profile of the cross-section of the nanoruler (*dashed line*) in the magnified inset, and its Gaussian fit (*continuous line*).

We next compared the performance of the *super*SIL microscope with that of a state-of-the-art, off-the-shelf STORM microscope (ZEISS Elyra equipped with an Alpha Plan-Apochromat 100×/1.46 NA oil immersion DIC objective), which we hereafter refer to as specialist STORM system. As a test sample, we imaged live *E. coli* cells expressing the antibacterial peptide ABC transporter McjD fused with enhanced GFP (EGFP) [29-32]. To reach quasi-equilibrium between the fluorescent on and off states of the EGFP molecules, power densities of 0.349 kW/cm^2^ and 1.1 kW/cm^2^ (λ = 488 nm) were delivered at the sample plane in the *super*SIL and specialist systems, respectively. As a control we imaged samples of *E. coli* expressing non-fluorescent McjD transporter with no EGFP (**Fig. S5**).

Typical FM images of EGFP-stained bacterial features from the *super*SIL and specialist microscopes are shown in **Fig. 2B**. Of note, the combination of a *super*SIL with a 100× dry objective lens results in a magnification of 471×, which is an enhancement of nearly 5 times over a conventional ZEISS 100× oil immersion objective. **Fig. 2C** shows representative STORM images and the corresponding molecular localization precision histograms. These reveal a 2-fold enhancement in localization precision using the *super*SIL STORM system (15.4 ± 4.1 nm) compared with the specialist STORM (31.3 ± 9.3 nm). *Super*SIL STORM also reported sub-cellular structures that the specialist STORM system failed to resolve because overlapping blinking events were not accounted for during the image reconstruction in order to achieve high accuracy of localizations. This illustrates the benefit of the increased magnification provided by the *super*SIL system in avoiding PSF crowding from high-density labelling scenarios, in particular when imaging smaller cells (e.g. bacteria, yeast, etc), in which it is difficult to resolve structural details [33].

Sample structure and labelling density can have an effect on resolution [34]. To ascertain the best molecular localization precision we could obtain with the specialist system, we imaged standard calibration samples of DNA origami [35]. These custom nanorulers (GattaQuant) carried ATTO 647N dye molecules, ~20 times brighter than EGFP [36]. As shown in **Fig. 2D**, results revealed a molecular localization precision of 17.6 ± 8.1 nm, in line with the microscope manufacturer’s specifications. Interestingly, the best localization precision we could extract from the specialist system using ATTO 647N was no better than the 15.4 ± 4.1 nm returned by the *super*SIL system using the much dimmer EGFP fluorophore. We therefore conclude that the *super*SIL microscope outperforms the specialist STORM system. Moreover, the *super*SIL microscope costs ~20 times less than the specialist STORM system.

We next characterized the performance of the *super*SIL system under cryogenic conditions (**Fig. 3**). We first verified that the freezing procedure does not alter the optical properties of the *super*SILs (**Fig. S6**). As shown in **Fig. 3A** and **3C**, the statistical results of PSF measurements from cubic zirconia defect spots confirmed that the resolution of the *super*SIL FM system (153 ± 14 nm) is not perturbed by being plunge-frozen together with the sample.

**Fig. 3.**
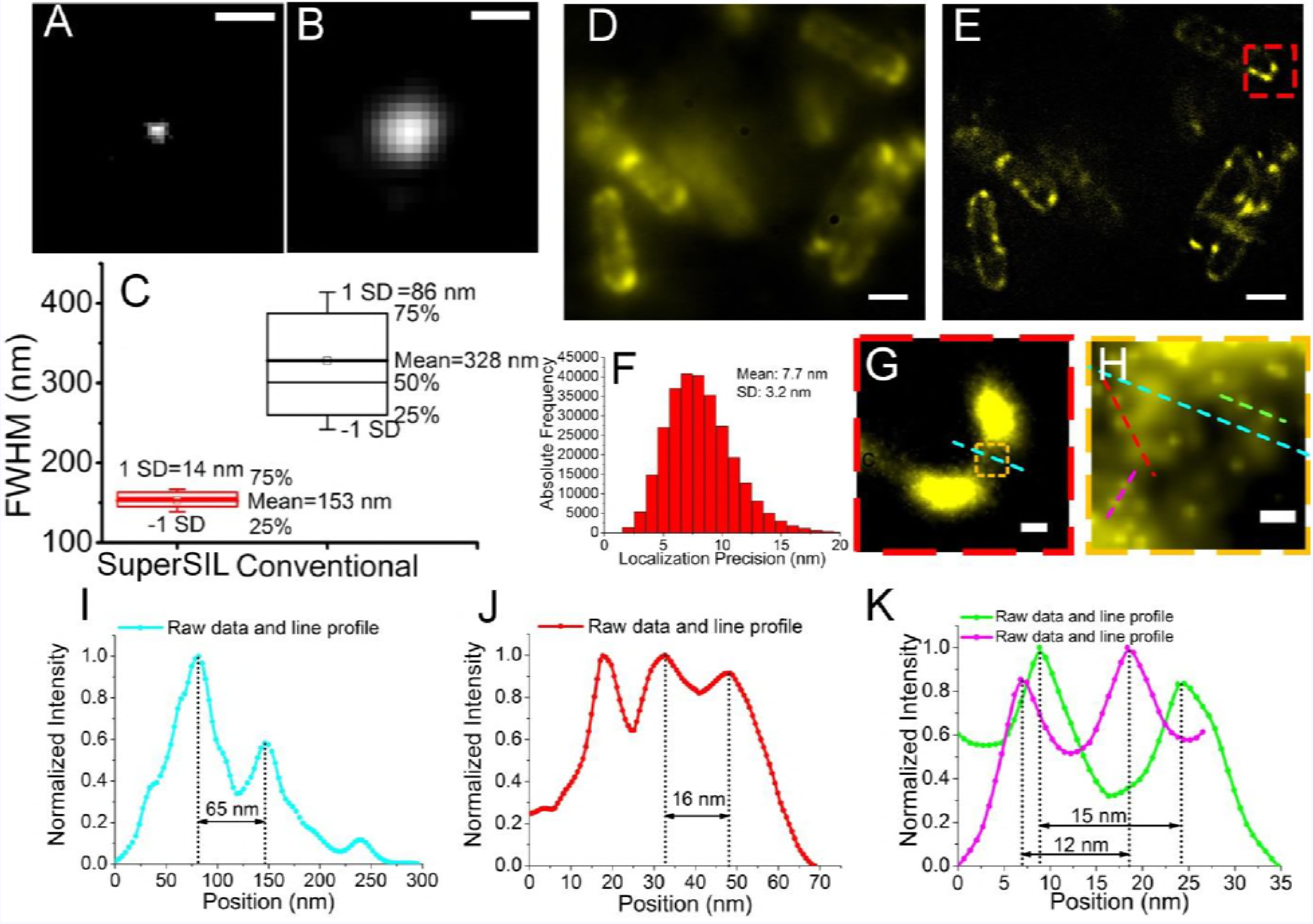
S*uper*SIL resolution performance under cryogenic conditions. Representative images of sub-diffraction limited objects from **(A)** the *super*SIL microscope and **(B)** the specialist system. Bar: 0.5 μm. **(C)** Box charts of FWHM of the PSFs for the *super*SIL and specialist systems showing the 25th and 75th percentiles of the data points. The whiskers show the standard deviation. **(D)** ATP-binding cassette (ABC) transporter protein PH1735 fused with EGFP in *E.coli* cells imaged in wide-field *super*SIL FM mode and **(E)** *super*SIL STORM mode. **(F)** Localization precision histogram from the image in **(E)**. **(G)** The enlarged image of the region of the cell indicated by the red dashed box in **(E)**. **(H)** The further enlarged image of the small region in the cell indicated by the orange dashed box in **(G)**. **(I)** Line profile of the cross-section of two PH1735 protein clusters indicated by the cyan dashed lines in **(H)**. **(J)** Line profile of the cross-section of two PH1735 protein clusters closer to each other indicated by the red dashed line in **(H)**. **(K)** Line profiles of the cross-section of two adjacent single molecules (magenta and green) indicated by the magenta and green dashed lines in **(H)**. Bars: 1 μm **(D, E)**, 100 nm **(G)** and 20 nm **(H)**.

For comparison, we evaluated the size of the PSF of a cryo-compatible FEI CorrSight fluorescence microscope equipped with a standard 0.9 NA 40x dry objective (ZEISS EC Plan-Neofluar 40×/0.9 Pol M27) and its auxiliary cryo-chamber, hereafter referred to as specialist cryo-STORM system. A representative image and the statistical results of the PSF measurement using sub-diffraction limited 100 nm fluorescent beads (ThermoFisher, TetraSpeck) are shown in **Fig. 3B** and **3C**. These results report a mean PSF size of (328 ± 86) nm, which is consistent with the theoretical value of 290 nm at the wavelength of 515 nm, according to Houston criterion [28]. Thus, the lateral resolution of *super*SIL FM is about 2-fold better than that of the specialist system.

We next characterized the resolution achievable using STORM. For this we imaged in LN_2_ vapour, plunged-frozen *E. coli* cells expressing the ATP-binding cassette (ABC) transporter PH1735, a putative multidrug transporter [37], fused to EGFP, which displayed robust single molecule blinking behaviour at cryogenic temperature (**Fig. S7A, S7B**). Representative wide-field and STORM images are shown in **Fig. 3D** and **3E**. The localization precision of 7.7 ± 3.2 nm (**Fig. 3F)** obtained with cryo-s*uper*SIL is 4.5-fold better than that of the specialist cryo-STORM system (35.7 ± 9.4 nm) (**Fig. S7C, S7D**). The latter value is consistent to that previously achieved in comparable conditions [10].

To reach quasi-equilibrium between the fluorescent on and off states of the EGFP molecules, we delivered a 1.1 kW/cm^2^ laser power at 488 nm wavelength to the sample plane of both the *super*SIL and specialist cryo-STORM systems. Interestingly, the substantially larger photon delivery and collection efficiency of the *super*SILs allowed us to use a 5.55 mW laser beam, whilst for specialist cryo-STORM imaging we required a 60 mW 488 nm laser beam. Thus, cryo-*super*SIL not only unlocks the potential to perform nanoscale STORM under cryogenic conditions, but makes this compatible with the use of low-power, low-cost lasers.

To illustrate the resolution of the cryo-*super*SIL system, we expanded the area within the red box in **Fig. 3E** (**Fig. 3G)**, and then further enhanced in the latter the area within the orange box. The resulting image (**Fig. 3H**) shows single PH1735-GFP molecules embedded in the membrane together with larger features. The profiles of fluorescence intensity across the areas marked in **Fig. 3H** reveal two features separated by 65 nm (**Fig. 3I**), two features separated by ~16 nm (**Fig. 3J**), and two pairs of molecules separated by 12 nm and 15 nm (**Fig. 3K**). PH1735 is a homodimeric ABC transporter (thus, fused with two EGFP molecules), and these features probably belong to a full transporter. Taking into account the linker length between the PH1735 and EGFP, the measured separation of 12-17 nm probably represents a nucleotide free inward-open transporter, which is consistent with distances measured on other ABC transporters such as McjD [31]. To our knowledge, the combination of the <8 nm localization precision and 12 nm resolution here demonstrated is the highest to date on cells. We objectively verified the resolution obtainable in cryo-*super*SIL STORM by imaging ATTO 647N-based DNA origami nanorulers (**Fig. S8**).

Two-colour imaging is required to study inter-molecular interactions. To investigate the possibility of two-colour imaging, we used McjD-EGFP expressing *E. coli* in which cell membranes were stained with the red fluorescent probe DiSC_3_(5). The DiSC_3_(5) dye is cationic and labels putative inner membrane nanodomains, i.e. the fluid lipid network [38], which is made of negatively charged phospholipids such as phosphatidylglycerol. In two-colour cryo-*super*SIL STORM imaging, laser beams of 1.1 kW/cm^2^ 488 nm and 2.2 kW/cm^2^ 642 nm powers were applied in the green and red channels. Two-channel imaging was implemented sequentially with the same camera setting, i.e. 30 ms exposure time per frame, 10 MHz 14 bit EM amplifier readout rate, 5.2× preamp setting and 300 EM gain. Channel alignment was implemented using feature detection during image post-processing. Representative images reveal the distribution of McjD proteins (*green*) and the organization of the fluid lipid network (*red*) (**Fig. 4A**), the latter displaying perceptible helical structures in agreement with previous findings [39]. Furthermore, a high degree of segregation is observed between the McjD and the lipid network. The localization precisions are 10.3 ± 3.8 nm in the green channel and 10.7 ±3.4 nm in the red channel (**Fig. 4B**).

**Fig. 4.**
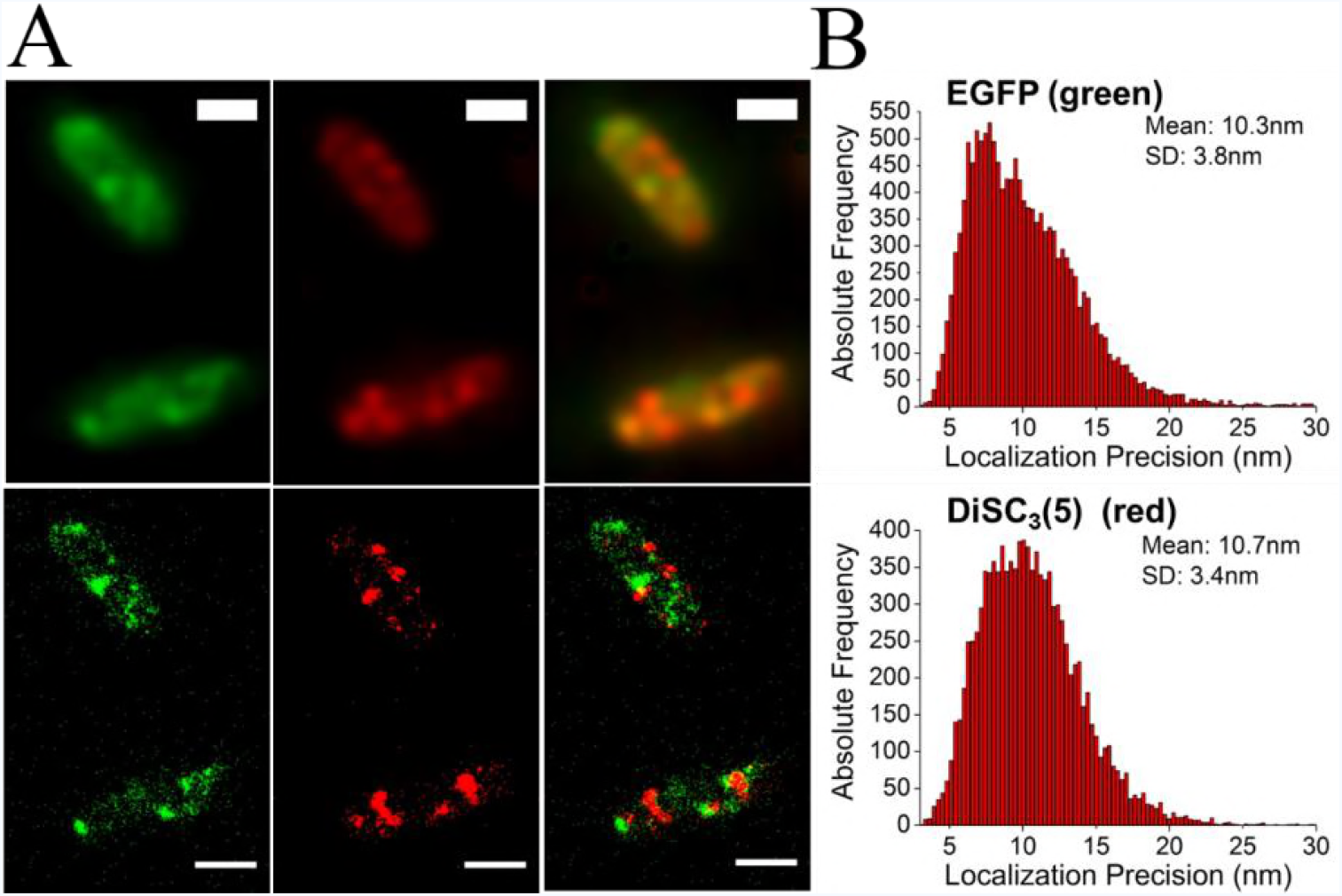
S*uper*SIL multi-colour imaging under cryogenic conditions. **(A)** McjD-EGFP (green) and membrane-DiSC_3_(5) (red) in *E.coli* cells imaged in (*top*) wide-field cryo-*super*SIL FM and (*bottom*) cryo-*super*SIL SMLM. The overlays of two-colour images are shown in the right-side column in each row. Scale bar: 1 μm. **(B)** The histograms of localization precision of (*top*) McjD-EGFP and (*bottom*) DiSC_3_(5).

Similarly to liquid immersion objective lenses, the depth of focus of the combination of the *super*SIL and the dry objective lens is inversely proportional to the square of the effective NA [27]. This results in a narrow depth of focus [40] and thus similar properties to TIRF. We found that the enhanced resolution of the *super*SIL is maintained up to separations of 100 nm from the flat surface of the lens (**Fig. S9-S10)**. This paves the way to deploy cryo-fixation for processes in mammalian cells that require TIRF. We have previously demonstrated TIRF using *super*SILs [21]. To verify that, under cryogenic conditions, the high resolution provided by the ultra-high NA = 2.17 *super*SIL is maintained in the basal periplasmic section adjacent to the lens surface, we imaged plunge-frozen Chinese hamster ovary (CHO) cells. In these the epidermal growth factor receptor (EGFR), a key molecule in cancer research [42], was labelled with Alexa Fluor 488. To extract the areas of highest resolution, we used a standard ‘rolling ball’ background reduction algorithm during image post-processing [43]. The TIRF-like appearance of the image is apparent in **Fig. 5A** [44]. A small cluster of EGFR in the cryo-*super*SIL FM image showed an apparent width of 230 nm (**Fig. 5B, 5C**). Two adjacent clusters 197 nm apart are also clearly distinguishable **(Fig. 5D, 5E)**. These values are indistinguishable to those predicted at depths from 10-100 nm (**Fig. S9**).

**Fig. 5.**
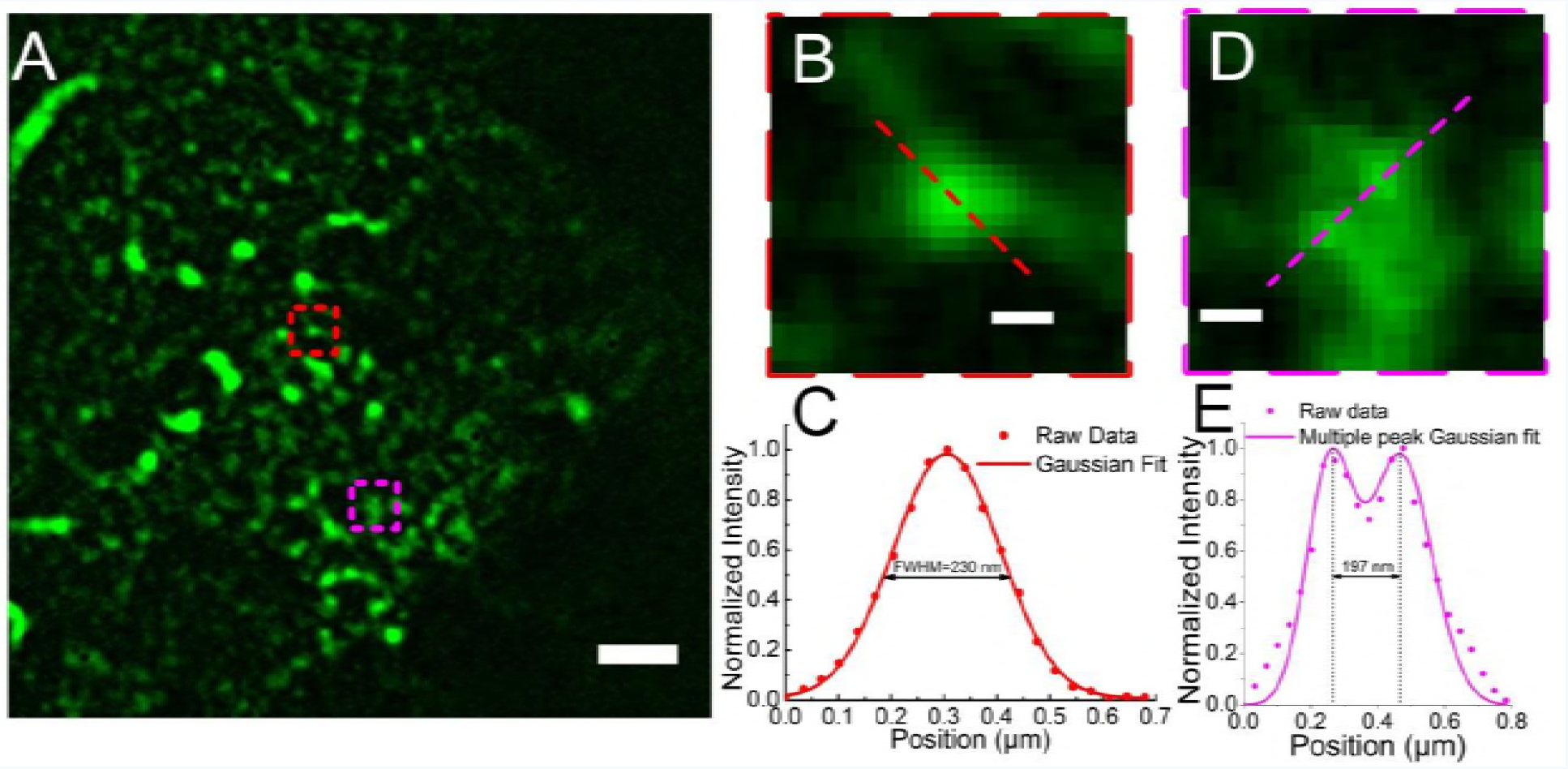
Wide-field *super*SIL FM at cryogenic temperature. **(A)** CHO cells expressing EGFR labelled with Alexa Fluor 488. **(B)** A cluster of EGFR indicated by the red dashed box in (B). **(C)** The line profile of the cross-section of the EGFR cluster (red dots) indicated by the red dashed line in (B), and its Gaussian fit (red curve). **(D)** Two adjacent EGFR clusters indicated by the magenta dashed box in (A). **(E)** The line profile of the cross-section of the two adjacent EGFR clusters (magenta dots) indicated by the magenta dashed line in (E), and their Gaussian fits (magenta curve). Bars: 2 μm (A) and 0.2 μm (B, D).

It is worth noting that the depth of focus of a *super*SIL microscope depends on the refractive index of the *super*SIL material, allowing a degree of depth “tuning”. This is an important consideration to exploit *super*SIL imaging in CLEM, where lamella thickness is in the range of 50- 300 nm [45]. If required, *super*SIL materials of lower refractive indexes (such as Quartz) can be used to increase the depth of focus approximately 2.2-fold (**Table S1**).

We therefore conclude that by combining a *super*SIL and a low NA dry objective we have achieved our goal of achieving super-resolution STORM in a conventional upright microscope. Indeed, the localization precision and resolution attained are, to our knowledge, the best obtained to date on cell samples. *Super*SIL-based super-resolution is not limited to STORM, but can be combined with other established super-resolution imaging techniques such as SIM [24] and STED [22]. Importantly, the high NA of the *super*SIL eliminates the barrier to combine cryo-imaging with TIRF, paving the way to the application of vitrification to super-resolution imaging also in mammalian cells.

During our characterization of the *super*SIL setup, we found an unexpected bonus. The ultra-high NA and increased magnification deliver a better resolution also at room temperature than that obtainable from an expensive, specialist STORM system. This means that, even if cryogenic conditions are not required, for example when imaging live cells, a simple add-on based around the use of *super*SILs would enable any non-expert with a basic fluorescence microscope to achieve state-of-the-art super-resolution at low cost. A simple modification of the *super*SIL assembly would easily enable the use of *super*SILs in inverted microscopes too. The method therefore overcomes the costs and complexity inherent to specialist super-resolution set-ups by circumventing the use of expensive objectives, intricate multi-stage illumination paths, specialized sample stages, and high power lasers. A comparison of *super*SIL microscopy vs fluid immersion microscopy is shown in **Table 1**.

**Table 1.** Characteristics of *super*SIL microscopy versus specialist fluid immersion microscopy.

| *SuperSIL* microscopy | liquid immersion microscopy |
| --- | --- |
| NAs in the range of 1.4-2.2, available as required | NA < 1.4 |
| Super-high resolution | High resolution |
| Super-high light collection efficiency | High light collection efficiency |
| Suitable for imaging under cryogenic conditions | Not suitable |
| Low cost | High cost |

Given these advantages, we propose that the low-cost, elementary *super*SIL technology has the potential to greatly extend the scope and the reach of super-resolution microscopy in cell biology. The availability to deliver close to 10 nm resolution under cryogenic conditions also has the potential to revolutionize cryo-CLEM, bridging the resolution gap between fluorescence and electron microscopy and allowing sample registration at the nanoscale. Possible schemes for CLEM include imaging of cryogenically-sectioned samples by cryo-*super*SIL super-resolution and transmission EM, or the use of focus ion beam scanning EM to image and produce lamellae suitable for the cryo-*super*SIL microscope. Given the possibility of scaling down *super*SILs to very small and custom-variable sizes, it is possible to envisage these devices being incorporated into EM systems, allowing true correlative microscopy without the need to move the sample and relocate areas of interest.

In summary, the ability of *super*SILs to achieve a very high NA simply and inexpensively without the use of immersion oil has the potential not only to transform super-resolution microscopy and CLEM, but also to increase the resolution attainable via fluorescence in any non-specialist laboratory at low-cost.

## Methods

### *Super*SIL microscope

The light source was a laser beam combiner (Omicron, LightHUB), including 488 nm and 642 nm laser lines, and a 470 nm collimated LED (Thorlabs, M470L3-C5). For the single colour imaging of McjD-EGFP and PH1735-EGFP *E. coli* cells, the filter sets consisted of a 469/35 nm bandpass filter (Semrock, FF01-469/35-25) as excitation filter, a 484 nm beam splitter (Semrock, FF484-FDi0) as dichroic filter, and a 512/25 nm bandpass filter (Semrock FF01-512/25-25) as emission filter. For the two-colour imaging of McjD-EGFP and membrane-DiSC_3_(5) *E. coli* cells, the filter sets consisted of quad-edge dichroic beam splitter (Semrock, Di01-R405/488/543/635-25×36) as dichroic filter and quad-band bandpass filter (Semrock, FF01-446/523/600/677-25) as emission filter, also used in the DNA origami nanoruler measurement.

### *Super*SIL assembly

The 1 mm diameter *super*SILs (Knight Optical, UK Ltd) were made of cubic zirconia (ZrO_2_), the cubic crystalline form of zirconium dioxide (ZrO_2_). The lenses have a refractive index of 2.17 and an Abbe number of 33.54 at the wavelength of 512 nm. This provides a high refractive index with medium dispersion suitable for *super*SIL FM. The assemblies were characterized by use of a coordinate measuring machine (OGP SmartScope ZIP 250 Coordinate Measuring Machine) to ensure the angle between the platinum disk and the *super*SIL’s flat surface is < 1°.

### SMLM data analysis

ZEISS ZEN software was used to process and render STORM images. In STORM image processing, various peak mask sizes were applied depending on pixel resolution and PSF size in each raw data set. The fit model was a two-dimensional Gaussian fit, and only single emitters from fluorophores were taken into account, whereas all multiple emitters were discarded. Following localization, displacements of molecules from drifts in the reconstructed images were corrected using feature detection and cross correlation. The counts from the raw images were firstly converted to the signal counts by deducting bias offset. Then the signal counts were converted to signal electrons by multiplying the preamp. Finally, the signal electrons were converted to photon numbers by adding the detector electron-multiplying gain.

### Sample preparation

#### Bacterial cell culture and staining

*E.coli* strain C43 expressing McjD fused with EGFP were prepared for imaging. Briefly, the overnight starter culture was diluted 1:100 in fresh LB media supplemented with 50 μg/ml kanamycin and grown at 37 °C until an optical density at 600 nm (O.D._600_) of 0.6 was achieved. Then the expression of McjD was induced by adding IPTG (Isopropyl β-D-1-thiogalactopyranoside) to a final concentration of 1 mM at 25 °C overnight. The cells were spun down before freezing. The EGFP counts were measured using a Spectramax microplate reader (Molecular Devices) from 1 ml of culture re-suspended in 200 μl PBS (Thermofisher). For the analysis, the bacterial pellet was re-suspended in 2 ml of PBS to an optical density (O.D._600_) of ~ 0.6 and kept on ice until plunge-freezing. For two-colour imaging, *E.coli* expressing McjD-EGFP were re-suspended in 250 μl PBS to O.D._600_ ~ 0.6 and centrifuged at 10000g for 5 min to wash out residues of culture medium. The pellet was then re-suspended in 25 μl of 100 nM DiSC_3_(5) in DPBS and incubated for 15 min on ice. 2.5 μl of sample were applied to the flat surface of each *super*SIL immediately prior to plunge-freezing.

#### Mammalian cell culture and staining

All reagents unless otherwise stated were from Thermo Scientific, UK. CHO cells expressing wtEGFR under an inducible Tet-ON promoter [46] were grown in 5% CO2 in air at 37 °C in phenol-red free Dulbecco’s Modified Eagle medium (DMEM) supplemented with 10% (v/v) fetal bovine serum, 2 mM glutamine, 100 μg/ml hygromycin B and 100 μg/ml geneticin. All cells used were regularly tested for mycoplasma contamination. Cells were seeded at a density of 10^5^/ml on superSILS passivated with PEG-BSA nanogel as described previously [47, 48]. Briefly, superSILs were etched with Piranha solution for 10 min, and thoroughly rinsed. Priming was performed for 5 min with Vectabond reagent (Vectorlabs) diluted 1:50 in acetone, followed by thorough rinsing. PEG-BSA nanogel was applied for 1h at 37 °C, rinsed twice with PBS, capped with 20 mg/ml BSA in PBS for 1 h 37 °C, quenched with 1 M Tris pH 8.0 for 15 min, followed by three washes in PBS. Cells were cultured for 48 h, rinsed, subjected to nutrient starvation for 2 h at 37 °C to wash out EGFR ligands from the serum and then labelled with 5 nM EGF conjugated to Alexa Fluor 488 (Thermo Scientific) for 30 min at 4 °C. Clustering of EGFR was induced by a 5 min incubation at 37 °C prior to plunge-freezing.

#### DNA origami nanorulers

Samples were prepared according to manufacturer’s instructions. Briefly, superSILs were glow discharged for 120 sec (negative) using a Quorum Q150T ES system, then washed 3 times with PBS, then immersed for 5 min in a solution of BSA-biotin 1 mg/ml in PBS and washed a further 3 times with PBS. Coating with neutravidin 1 mg/ml in PBS was also performed by immersion for 5 min, followed by three washes in PBS + 10 mM MgCl_2_ (immobilization buffer). 2.5 μl of DNA Origami diluted in immobilization buffer to a final concentration of 1:100 were applied to the flat surface of the *super*SILs, prior to blotting and plunge-freezing.

#### Plunge freezing

The superSILs were etched for 15 min with Piranha solution (3:1 concentrated sulphuric acid, 30% H_2_O_2_, both from Sigma-Aldrich) then rinsed with plenty of water and left to air dry for 1 h. SuperSILs were then glow-discharged for 120 s (negative) prior to sample loading and freezing using a Quorum Q150T ES system. Samples were frozen by plunge-freezing using FEI Vitrobot MKIV according to manufacturer’s instructions. The chamber was equilibrated to 4 °C, 95% relative humidity. Blotting was performed manually.

## Author Contributions

* L.W. and M.L.M conceived and designed the research. B.B. and L.W. constructed the cryo-SIL microscope and implemented the measurements. S.A. and C.S. manufactured the superSIL assembly. L.C.Z. and M.C.D. developed sample plunge freezing protocol. L.C.Z., M.R., A.N.M and S.R.N. prepared biological samples. K.B. supervised M.R. in the bacterial sample preparation. L.W., B.B., D.R., D.T.C and M.L.M analysed data. D.T.C., M.L.M and L.W. wrote the manuscript. All the authors discussed and commented on the manuscript.

## ACKNOWLEDGEMENTS

We thank John Collier for his continuous support and contributions to this project and Simon Nestead and David Drew for kindly providing us with the expression construct for PH1735. This work has been funded by Medical Research Council grants (MR/K015591/1) to MMF and (MR/N020103/1) to KB.

